# A purely visual adaptation to motion can differentiate between perceptual timing and interval timing

**DOI:** 10.1101/2023.02.20.529202

**Authors:** Aurelio Bruno, Federico G. Segala, Daniel H. Baker

**Affiliations:** Department of Psychology, University of York, Heslington, York, YO10 5DD, UK; School of Psychology and Vision Sciences, College of Life Sciences, University of Leicester, University Road, Leicester, LE1 7RH, UK

**Author notes:** Corresponding author: Aurelio Bruno.

**Keywords:** time perception, duration, adaptation, computational modelling, time scales

## Abstract

It is unclear whether our brain extracts and processes time information using a single centralized mechanism or through a network of distributed mechanisms, which are specific for modality and time range. Visual adaptation has previously been used to investigate the mechanisms underlying time perception for millisecond intervals. Here, we investigated whether a well-known duration aftereffect induced by motion adaptation in the sub-second range (referred to as ‘perceptual timing’), also occurs in the supra-second range (called ‘interval timing’), which is more accessible to cognitive control. Participants judged the relative duration of two intervals after spatially localized adaptation to drifting motion. Adaptation substantially compressed the apparent duration of a 600 ms stimulus in the adapted location, whereas it had a much weaker effect on a 1200 ms interval. Discrimination thresholds after adaptation improved slightly relative to baseline, implying that the duration effect cannot be ascribed to changes in attention or to noisier estimates. A novel computational model of duration perception can explain these results, and also bidirectional shifts of perceived duration after adaptation reported in other studies. We suggest that we can use adaptation to visual motion as a tool to investigate the mechanisms underlying time perception at different time scales.

## Introduction

The ability to code and process time information in the millisecond range is essential for several everyday activities, ranging from action coordination, in the motor domain, to speech processing and recognition, in the language domain, to motion detection and processing, in the sensory domain, and to more sophisticated behaviours like social interactions mediated by gaze. Although temporal processing on this scale is “[…] *probably the most sophisticated and complex form of temporal processing*” (Mauk & Buonomano, 2004), our knowledge of the underlying brain mechanisms remains quite poor. A similar lack of certainty affects the study of time perception in the order of seconds, where time is estimated in a more conscious fashion.

Some theories propose a single centralized mechanism, an ‘internal clock’, which would encode the duration of any interval regardless of the sensory modality of the embedded sensory stimulus, and of the time scale of the interval itself (Creelman, 1962; Treisman, 1963; Treisman et al., 1990), simply by integrating the number of pulse-like signals generated by a pacemaker between the onset and offset of the considered interval. Empirical support for this model comes from the observations that increased arousal or attention induce duration overestimation, as they would speed up the clock (Droit-Volet & Wearden, 2002; Wearden et al., 1999), and that we cannot time two events simultaneously (Morgan et al., 2008). No biological substrate for this single time mechanism has been identified yet. Other theories suggest that time perception might be the product of a network of distributed mechanisms, which are modality-specific and contribute independently to time processing according to the time scale of the involved intervals (Bruno & Cicchini, 2016). Empirical support for this idea comes from the observations that many time perception biases like central tendency effects (Murai & Yotsumoto, 2016), rate aftereffects (Motala et al., 2018) or duration aftereffects (Heron et al., 2012) show little or no cross-modal transfer. Evidence exists of distributed mechanisms at both cortical and subcortical levels (Binetti et al., 2020; Heron et al., 2019; Paton & Buonomano, 2018).

Time perception in the sub-second range has been often referred to as ‘perceptual timing’ for being more tightly linked to perceptual processing and less susceptible to the influence of cognitive control, whereas, in the supra-second range, we refer to it as time estimation or ‘interval timing’, which is under a more direct control of higher cognitive functions like memory (Michon, 1985). The idea that this distinction is reflected in different underlying mechanisms in our brain has received empirical support from psychophysical (Murai & Yotsumoto, 2016; Rammsayer et al., 2015), neuroimaging (Lewis & Miall, 2003a, 2003b; Nani et al., 2019; Wiener et al., 2010), neurophysiological (Kunimatsu et al., 2018) and pharmacological (Rammsayer, 2008) studies.

Over the last few years, adaptation has been extensively used as a tool to study the mechanisms underlying time perception. At least two different types of adaptation were shown to induces changes in the perceived duration of sub-second intervals. First, adapting to several repetitions of an interval with a long or short duration resulted in repulsive duration after-effects, the magnitude of which depended on the temporal distance between the adaptor and test lengths (Heron et al., 2012; Walker et al., 1981) - but see Curran et al. (Curran et al., 2016). These observations suggested the existence of duration-selective mechanisms with similar characteristics to those which process the spatial frequency or the orientation of a visual stimulus: adapting a given duration channel shifts the peak of the neuronal population response away from that duration, influencing the subsequent time judgments accordingly. There is a second type of adaptation, which does not address a specific duration channel and, nonetheless, induces a change in the apparent duration of a subsequent interval. A purely visual adaptation to motion or flicker can in fact produce a substantial perceptual compression of duration, which is temporal-frequency dependent, and it is limited to the adapting location (Burr et al., 2007; Gulhan & Ayhan, 2019; Johnston et al., 2006). The mechanisms involved in this type of adaptation are arguably not specific to the processing of duration, and they are most likely responsible for the processing of visual motion and temporal change.

While there is some evidence suggesting that the effect of duration adaptation might extend to supra-second intervals (Shima et al., 2016), we do not know yet whether adaptation to visual motion shows the same flexibility. If there was a dissociation between the two time scales in the adaptation-induced duration changes, that would not only provide further evidence supporting the existence of time-scale dependent channels of time perception, but it would also shed some light on the brain sites where this type of adaptation takes place, which are still unknown (Burr et al., 2011; Johnston et al., 2011).

In order to address this issue, in this study, we measured perceived duration after space-specific adaptation to drifting motion for intervals centred around a sub-second duration (600 ms) or a supra-second duration (1200 ms), in separate sessions. Furthermore, we investigated whether adaptation interfered with our participants’ ability to discriminate changes in duration and whether changes in perceived duration depend on changes in duration discrimination after adaptation in the two time ranges. Finally, we developed a computational model of duration perception that can account for both duration compression and bidirectional repulsive duration aftereffects.

## Methods

### Observers

Twenty observers (including two authors) participated in the experiment. All of them had normal or corrected-to-normal vision. All observers provided written informed consent prior to testing, and procedures were approved by the ethics committee of the Department of Psychology at the University of York (Ethics Application ID: 782).

### Apparatus

Stimuli were displayed, in a dark room, on a gamma-corrected LCD monitor (ASUS ROG Swift PG258Q), with a refresh rate of 240 Hz. We confirmed that our monitor indeed provided the correct timing of visual stimulus presentation by measuring different stimulus durations with a photodiode measurement circuit connected to an oscilloscope. The stimuli were generated in Matlab using the Psychophysics Toolbox extensions (Brainard, 1997; Pelli, 1997). Stimuli were viewed from a distance of 57 cm. The head of participants was restrained with a chinrest.

### Procedure

All participants completed, in different sessions, an adaptation condition for a sub-second duration (600 ms), an adaptation condition for a supra-second duration (1200 ms) and two baseline conditions, one for each duration. In the baseline conditions, no adaptation was presented, and participants always completed them before the adaptation conditions. The adaptation experiment was composed of an adaptation phase followed by a test phase (Figure 1). Continuous fixation on a central spot was required for the whole duration of the experiment. All stimuli were drifting luminance-modulated Gabors (vertically oriented, spatial frequency: 1 cycle/degree,) displayed at a distance of 5° of visual angle from the centre of the monitor. The diameter of the stimulus window was 5° of visual angle, and the Standard Deviation of the Gaussian spatial envelope was 0.83° of visual angle. Michelson contrast was 50% for the adaptor and 80% for the tests to avoid reductions in apparent contrast in the tests after adaptation (Georgeson, 1985). In the adaptation phase, participants were exposed to an eccentric adaptor (displayed on the left-hand side of the screen, 5° from the monitor centre), which reversed direction every 500 ms to avoid inducing a directional motion after-effect. The total adaptation time was 32 s for the first trial (8 s for the following trials), divided into eight cycles of 4 s each (four cycles of 2 s each for the following trials). In each cycle, the adapting speed could be either 5 or 20 ° /s (the presentation order of the cycles was randomly interleaved on a trial-by-trial basis) so that the proportion of 5/20 ° /s adaptation was 50%-50%. We chose this particular proportion as it was shown to minimize the effect of adaptation on the perceived speed of the tests (Ayhan et al., 2011; Ayhan et al., 2009; Bruno et al., 2010). When the adaptor disappeared, it was replaced by a blank screen of mean luminance for 500 ms, which preceded the test phase. In the test phase, two test stimuli were sequentially displayed: one, the standard, in the same location as the adaptor and the other, the comparison, in the opposite position relative to the fixation spot. The presentation order was randomized, and the two test intervals were separated by a 500-ms uniform mean luminance screen. Both tests drifted at 10 ° /s in opposite directions relative to each other. The standard had fixed duration across trials (either 600 or 1200 ms, in separate sessions), whereas the duration of the comparison varied in seven fractions (i.e., 1/3, 2/3, 5/6, 1, 7/6, 4/3, 5/3) of the standard duration in order to generate a psychometric function (for each comparison duration, we ran at least 20 repetitions). After the disappearance of the second test, participants were required to report which test had stayed on for the longer duration, by pressing one of two designated response buttons on a computer keyboard. We used custom Matlab code to fit cumulative Gaussian functions through the individual and mean data. The Point of Subjective Equality (PSE, defined as the 50% point on the psychometric function) was our measure of perceived duration, whereas the Just Noticeable Difference (JND, defined as half the difference between the 75% and the 25% points on the psychometric function) was our measure of duration discrimination.

**Figure 1.**
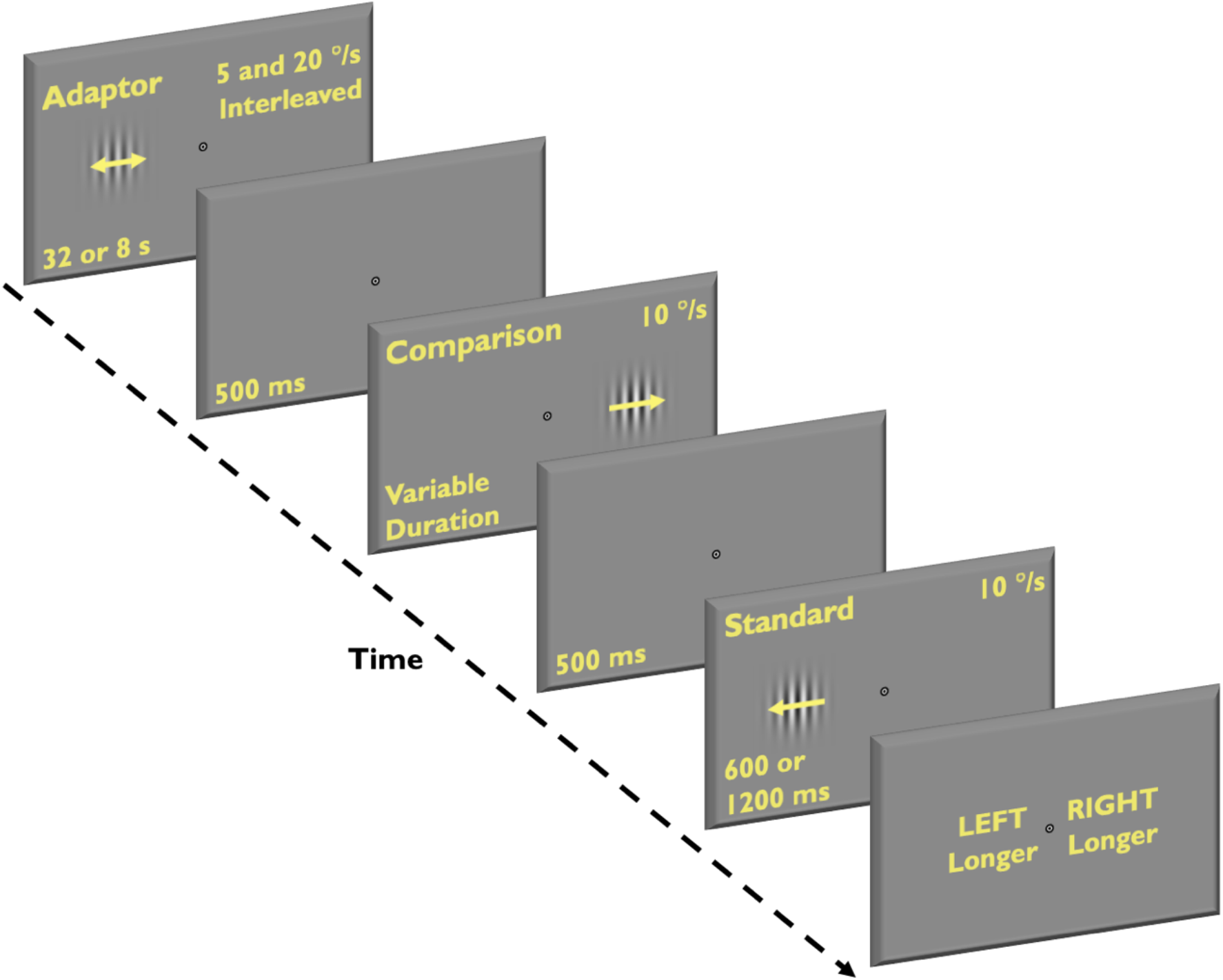
Schematic illustration of the procedure adopted in the adaptation conditions of the experiment. Participants adapted to an oscillating Gabor stimulus, which was displayed in a specific spatial location on the screen. The adaptor drifted at a speed, which varied in equal cycles of 5 and 20 ° /s (interleaved within the same adaptation period to minimize the effect on the perceived speed of the following tests). Two test stimuli (also Gabors, drifting at 10 °/s) were subsequently displayed, one after the other: the standard, in the same location previously occupied by the adaptor; the comparison, in an unadapted location. Participants were required to report which test had the longer duration.

## Results

We measured participants’ ability to estimate the relative duration of two intervals containing Gabor stimuli drifting at 10 ° /s, displayed in the near periphery of a central fixation, after adaptation to visual drifting motion (of a combination of 5 and 20 ° /s speed in equal proportions to minimize the effect of the adapting speed on that of the test stimuli). The duration of the comparison stimulus varied, on a trial-by-trial basis, relative to that of the standard, which remained constant. The resulting psychometric functions (averaged across twenty participants) are plotted in Figure 2 for the baseline and adaptation conditions and for the sub-second (600 ms, Figure 2a) and supra-second (1200 ms, Figure 2b) durations. We considered the duration, in correspondence of which participants’ performance was at chance (i.e., the Point of Subjective Equality, PSE), as a measure of the subjective duration of the standard interval. Durations are expressed as percentages of the standard duration to facilitate the comparisons between physically different durations. The data represented in Figure 2 are averaged across all participants, however perceived duration estimates, and duration discrimination estimates described below were derived from individual fits.

**Figure 2.**
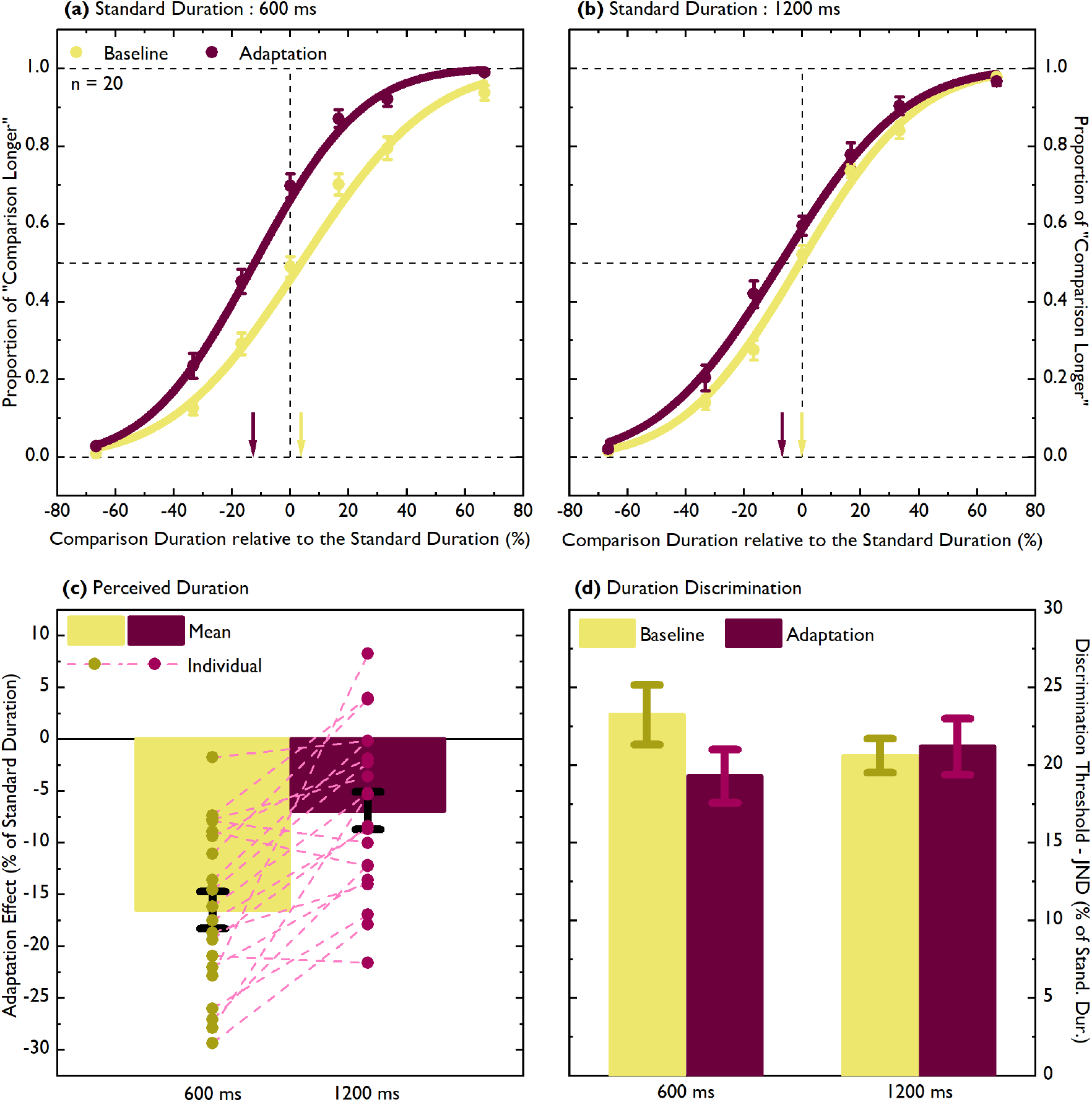
(a) Mean psychometric functions (averaged across twenty participants) are plotted for the baseline (yellow circles and curve) and the adaptation conditions (dark red circles and curve), for the sub-second duration (600 ms) The proportion of “Comparison Longer” responses is plotted as a function of the comparison stimulus actual duration, expressed as a percentage change relative to the standard stimulus duration: “0” represents the actual duration of the standard stimulus, positive values indicate comparison durations longer than the standard duration, negative values comparison duration shorter than the standard duration. The dark red and yellow arrows represent the Points of Subjective Equality for the adaptation and baseline conditions, respectively. Negative PSEs indicate duration compression. (b) Same as in (a) for the supra-second duration (1200 ms). (c) The mean (columns) and individual (circles) adaptation effects, defined as the difference between the PSE obtained in the adaptation condition and the PSE obtained in the baseline condition expressed as a percentage of the standard duration, are plotted for 600 (yellow) and 1200 (dark red) ms. The dashed lines connect each participant’s effects in the two standard duration conditions. (d) The Just Noticeable Differences are plotted for the baseline and adaptation conditions, and for 600 and 1200 ms standard durations. Error bars represent ± 1 Standard Error Mean (S.E.M.).

Overall, in the baseline conditions of both standard durations, our participants performed the task accurately: short comparison durations were infrequently judged as being longer than the standard duration, whereas for long comparison durations, the proportion of “longer” responses was higher than chance. When standard and comparison intervals had the same duration (i.e., the “0” duration in the plots), participants’ performance was at chance (even though, at 600 ms, the PSE was significantly higher than 0, PSE Baseline 600 ms = + 3.74 %, Standard Error Mean = 1.53, one-sample *t* test, *t*(19) = 2.45, *p* = 0.024). For the sub-second duration (Figure 2a) adaptation induced a substantial leftward shift in the psychometric function relative to the baseline, which indicates a subjective compression of duration. The PSE after adaptation was significantly shorter than in the baseline condition (PSE Adaptation 600 ms = -12.75 %, S.E.M. = 1.8, paired-samples *t* test, *t*(19) = 9.27, *p* < 0.0001). This result confirms previous observations obtained after adaptation to visual motion or flicker in the millisecond range (Bruno & Cicchini, 2016; Burr et al., 2007; Johnston et al., 2006). The effect of this type of adaptation for longer durations had not been systematically investigated yet. For our supra-second duration (1200 ms, Figure 2b), the duration compression observed after adaptation (indicated by the leftward shift of the psychometric function) was substantially less pronounced, though still significant (PSE Baseline 1200 ms = - 0.127, S.E.M. = 1.08; PSE Adaptation 1200 ms = - 7.02 %, S.E.M. = 1.93; paired-samples *t* test, *t*(19) = 3.843, *p* = 0.001). Overall, adaptation induced a significant duration underestimation relative to a condition without adaptation (ANOVA Repeated Measures, Main Effect Adaptation, *F* (1,19) = 57.71, *p* < 0.0001), but the effect depended on the standard duration (Interaction Adaptation x Standard Duration, *F*(1,19) = 27.92, *p* < 0.0001).

In order to compare the relative magnitude of the adaptation-induced changes in perceived duration for our sub-and supra-second intervals, in Figure 2c, we plotted the adaptation effect, defined as the difference between adaptation and baseline estimates, expressed as percentage of the standard duration. The adaptation effect for a 600 ms standard duration was more than twice as large as that for a 1200 ms duration (Adaptation Effect 600 ms = -16.49 %, S.E.M. = 1.78; Adaptation Effect 1200 ms = -6.89 %, S.E.M. = 1.79; paired-samples *t* test, *t*(19) = -5.284, *p* < 0.0001). Each dashed line in Figure 2c connects the individual adaptation effect for 600 ms with the corresponding adaptation effect for 1200 ms for the same participant. It can be noticed that, first, seventeen out of twenty participants showed a stronger adaptation-induced duration compression for the sub-second duration and, second, that most participants showed a remarkably similar difference between the two durations (as highlighted by the fact that most of the dashed lines are almost parallel), indicating that the observed group effects well captured the individual patterns of results.

The duration compression described thus far could arguably be a consequence of a reduced sensitivity to duration changes (i.e., noisier estimates) after adaptation. If a change in the PSE was accompanied by a corresponding change in the slope of the psychometric function after adaptation, which would indicate a change in duration discrimination, it would be hard to claim that adaptation induced a specific effect on the subjective estimates of duration. Following the same logic, the difference in the magnitude of the adaptation effect between sub- and supra-second durations could potentially be due to different changes in duration discrimination in the two time scales. From Figure 2, we can tell at a glance that this scenario appears to be unlikely in our data: adaptation does not seem to dramatically change the slopes of the psychometric functions. A statistical analysis confirmed this impression. In Figure 2d, we plotted the Just Noticeable Difference (defined as the difference between the 75% point and the 25% point on the psychometric function, divided by two), as a measure of duration discrimination threshold, for all the conditions in the study. Overall, duration discrimination improved after adaptation (ANOVA repeated measures, Main Effect Adaptation, *F*(1,19) = 5.016, *p* = 0.037). Even though the interaction between the adaptation and standard duration factors did not reach statistical significance (*F*(1,19) = 3.985, *p* = 0.06), direct comparisons between adaptation and baseline conditions showed that the mean JND was lower after adaptation for 600 ms (JND Baseline 600 ms = 23.24 %, S.E.M. = 1.92; JND Adaptation 600 ms = 19.3 %, S.E.M. = 1.71; paired-samples *t* test,, *t*(19) = 2.892, *p* = 0.009) but not for 1200 ms.

In principle, there could still be a significant individual trend that linked changes in perceived duration to changes in duration discrimination. Group statistics cannot, in fact, completely exclude that a particular direction in the duration estimates (towards duration compression, for example) was systematically associated to noisier or better duration discrimination. In order to investigate this possibility, we plotted the individual PSEs as a function of the individual JNDs, for both 600 ms and 1200 ms standard duration (see Electronic Supplementary Material, Figure S1). The correlation between the two measures reached statistical significance only for the baseline condition at 600 ms (Pearson’s *r* = 0.669, *p* = 0.001), indicating that the higher the duration discrimination threshold were, the larger the overestimation of interval duration for this condition. We can therefore claim that changes in perceived duration after adaptation did not depend on changes in duration discrimination.

One might wonder whether the stronger adaptation effect observed for 600 ms relative to 1200 ms standard duration was in fact driven by differences already present in the baseline conditions for the two durations. As we reported above, mean estimates for 600 ms were found to be significantly higher than 0, indicating duration overestimation. Moreover, the amount of this overestimation was positively correlated with the duration discrimination thresholds (see Electronic Supplementary Material, Figure S1a). In order to probe this alternative explanation, we re-analyzed the data excluding the three participants with the highest PSEs for the baseline condition at 600 ms. They all hugely overestimated the standard duration in the absence of adaptation (by 11.07%, 15.73% and 21.73%, respectively), and their PSEs appear to be substantially separated from those of the rest of participants (see Electronic Supplementary Material, Figure S1a). This exclusion was enough to reduce the average PSE to 1.54 % (S.E.M. = 1.02), which was no longer significantly higher than 0. The correlation between the PSEs and the JNDs for the same condition also became non-significant. However, the adaptation effect for 600 ms remained twice as large as for 1200 ms (Mean Adaptation Effect 600 ms = - 15.66 %, S.E. M. = 2.03; Mean Adaptation Effect 1200 ms = - 6.97 %, S.E. = 1.82; paired-samples t-test, *t*(16) = -5.227, *p* < 0.0001), discarding the possibility that this difference was due to a tendency to overestimate duration without adaptation.

## Computational modelling

Adaptation aftereffects such as the tilt aftereffect are typically modelled using a population of tuned units that tile the stimulus space (Clifford et al., 2000). Perception of a given stimulus is determined either by the mechanism with the greatest response, or by a more sophisticated population read-out rule such as maximum likelihood decoding (Jazayeri & Movshon, 2006). Adapting to a particular stimulus shifts the peaks of the adjacent mechanisms away from the adaptor, producing a bidirectional repulsive aftereffect. This general scheme was proposed by Heron et al. (Heron et al., 2012) to account for their bidirectional duration aftereffects. However, their model is implemented only at the stage of tuned duration channels and lacks a front end that can process stimuli with arbitrary temporal properties (such as the drifting stimuli we use here).

Our aim was to construct a model that could account for both the repulsive aftereffects observed for a fixed-duration adaptor, and the duration compression effects we report here when using a drifting adaptor. To do this, we converted a model developed by Meese and Baker (Meese & Baker, 2023) to explain analogous aftereffects for stimulus size (i.e. a 2 deg adaptor causes 1 deg targets to appear smaller, and 4 deg targets to appear larger) for use in the time domain. A more extensive derivation of the model can be found in the original paper (Meese & Baker, 2023). In brief, incoming stimuli are processed by a bank of mechanisms sensitive to different durations (Figure 3a). The output of each mechanism is then subject to nonlinear transduction, which involves divisive suppression from the largest mechanism (in the spatial domain this is surround suppression). Across the population of mechanisms, the response to a single stimulus (examples in Figure 3b) is then a saturating sigmoidal function (black curve in Figure 3c). To determine the peak response, the derivative is calculated by taking the difference between each mechanism and its neighbour (blue curve in Figure 3c).

**Figure 3.**
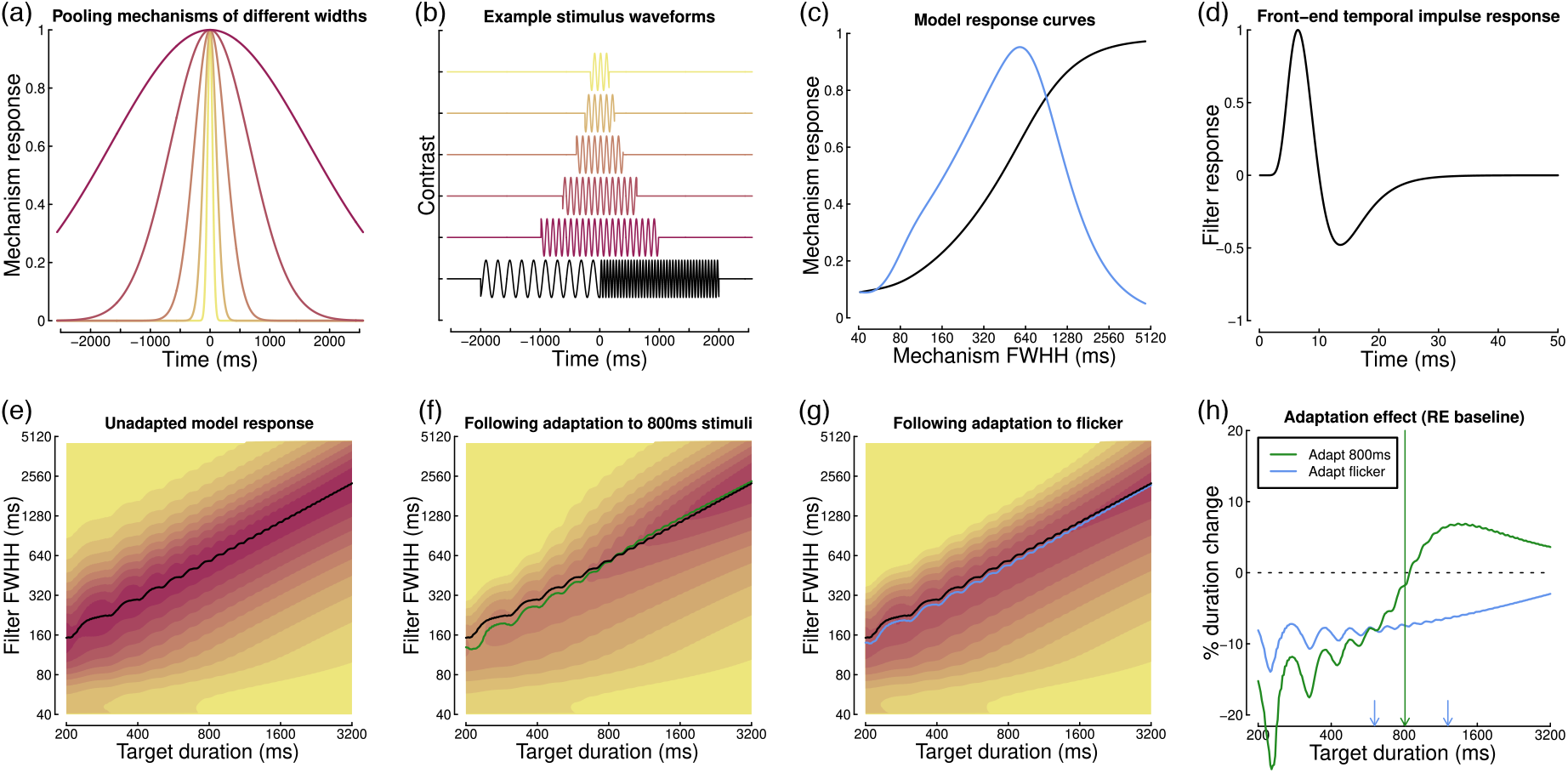
Summary of model components and behaviour. Panel (a) shows example pooling windows for five mechanisms that integrate over different durations. Panel (b) shows example stimulus waveforms for 10Hz modulated targets of different durations, and the flickering 5Hz/20Hz adaptor (black). Panel (c) shows model responses to an 800ms target, as a function of mechanism width. The black curve is the output of the nonlinear transducer, and the blue curve is the derivative (obtained via subtraction between adjacent mechanisms). Panel (d) shows the temporal impulse response function used as a front-end filter for the model. Panel (e) shows the model response as a joint function of target duration and filter width, with the black curve showing the location of the peak response prior to adaptation. Panels (f) and (g) show the model response following adaptation to either an 800ms 10Hz target (f) or the 5Hz/20Hz flicker (g). Black curves in panels (f,g) are duplicated from panel (e), and the green and blue curves indicate the peaks following adaptation. Panel (h) shows the predicted change in perceived duration after adapting to a fixed duration (800ms) or a flickering adaptor.

In order to handle arbitrary temporal waveforms, we added a ‘front end’ to the model, consisting of a temporally band-pass biphasic impulse response function (see Figure 3d). We did consider using multiple temporal filters, as suggested by Johnston (Johnston, 2010, 2014), however these were not required to reproduce our effects. Other filters, such as a low-pass filter, behaved similarly for the conditions we consider. Adaptation is implemented by adjusting the gain of each mechanism in proportion to its response to the adapting stimulus. See Meese and Baker (Meese & Baker, 2023) for a formal mathematical account, and the online modelling code for implementation.

Figure 3e illustrates that, prior to adaptation, the model gives a duration read-out (black curve) that increases in proportion to stimulus duration. Note that the ‘wiggles’ in the curve at brief durations are due to interactions between the phase of the sinusoidal carrier and the stimulus duration envelope. When the model is adapted to a single duration (800ms) perceived duration is shifted away from the adaptor duration in both directions (see the green curve in Figure 3f,h). Adapting to flicker at 5Hz and 20Hz (equivalent to a drifting stimulus from the perspective of a local one-dimensional filter) instead produces a unidirectional shift, whereby perceived durations of all target stimuli are reduced (blue curve in Figure 3g,h). However, this effect is largest at shorter durations, and reduces for longer durations, in qualitative agreement with our empirical results (see Figure 2c). The model architecture is therefore able to explain the main features of both repulsive and compressive aftereffects within a single framework.

## Discussion

We used a well-established adaptation paradigm (Bruno & Cicchini, 2016; Johnston et al., 2006) to investigate the effect of adaptation to visual motion on perceived duration and duration discrimination for both sub- and supra-second intervals. We found that adaptation perceptually compressed the duration of a 600 ms interval and that the magnitude of the effect relative to a baseline condition without adaptation corresponded to about 16% of the interval length. For a 1200 ms interval, we observed a much weaker duration compression after adaptation: the adaptation effect amounted to about 7% of the interval length. Duration discrimination improved slightly after adaptation, especially for the sub-second duration (i.e., 600 ms). We did not observe any significant relationship between the changes in perceived duration after adaptation and the corresponding changes in duration discrimination.

The distinction between ‘perceptual timing’ and ‘interval timing’ for the processing of sub- and supra-second intervals, respectively, was mainly based on the absence (for the former) or presence (for the latter) of cognitive control. Perceptual timing would rely on more sensory and automatic processes, which arguably occur in early pre-cortical and cortical sensory sites, whereas interval timing would depend on more cognitive processes, which arguably occur in higher-order cortical areas. Recent systematic meta-analyses of neuroimaging studies that used timing tasks confirmed these predictions. Even though there was a quite high degree of overlap between networks of areas involved in millisecond and second processing (mainly in the frontal lobes, where the Supplementary Motor Area was almost always active in relation to timing tasks), subcortical areas, like the cerebellum and basal ganglia, were found to be more involved in sub-second tasks, whereas supra-second tasks required more cortical contributions, especially from the inferior parietal cortex (Nani et al., 2019; Wiener et al., 2010). Studies using transcranial magnetic stimulation reported disruptions of the performance in timing tasks for sub-second, but not supra-second intervals after stimulation to the cerebellum and the opposite pattern when the stimulation occurred over the dorsolateral prefrontal cortex (Jones et al., 2004; Koch et al., 2007). The difference we observed in this study between the effect of adaptation on sub-second intervals relative to that on supra-second intervals might therefore suggest that this adaptation addressed more subcortical, rather than cortical components. This is consistent with previous observations that adapting to a cortically invisible temporal frequency, above the flicker fusion threshold, still induced a spatially localized duration compression (Johnston et al., 2008). In a recent EEG study, Tonoyan et al. looked at the electrophysiological signal in a perceived duration task in the sub-second range, after adaptation to visual motion (Tonoyan et al., 2022). In terms of event-related potentials, they observed that changes in the amplitude of the N200 component (which is thought to arise from area V5/MT), recorded in contralateral occipital electrodes, reflected changes in perceived duration after adaptation. They also found that changes in Beta power after adaptation could predict perceptual duration compression. Taken together, these results suggest that the effect of motion adaptation on perceived duration in the sub-second range occurs locally and relatively early in the visual system.

It has to be noted that the effect of adaptation on perceived duration of a 1200 ms interval was weak, but did not completely disappear in our data. This could be due to the fact that this duration is barely in the supra-second range and the shortest two of the seven comparison durations we used were in the sub-second range. More generally, it is not clear where the boundary between sub- and supra-second processing can be traced in terms of interval duration (Paton & Buonomano, 2018). We chose 1200 ms and not a longer duration to prevent our participants from using a counting strategy to estimate duration (Grondin et al., 1999).

It is worth stressing again that the adaptation to a period of continuous visual drift we used here is different from the adaptation to several repetitions of a specific duration described by Heron et al. (Heron et al., 2012), which we could call ‘duration adaptation’. In our case, in fact, we were not adapting a putative duration channel, but rather a channel sensitive to visual speed or temporal frequency. To avoid confusion, we will call it ‘motion adaptation’. We showed here that the effect of motion adaptation on duration perception is significantly weaker in the supra-second scale. On the contrary, Shima et al. reported comparable effects after duration adaptation for the two time ranges and that adaptation transferred from one range to the other (Shima et al., 2016). This difference is hardly surprising, as duration adaptation and motion adaptation were shown to differ across other dimensions as well. Duration adaptation in the visual domain, in fact, shows a very broad spatial tuning (Fulcher et al., 2016; Li et al., 2015; Maarseveen et al., 2017) and a strong interocular transfer (Heron et al., 2019), suggesting a late rather than early cortical site for this kind of adaptation to occur. Consistent with this prediction, neuronal activity in the inferior parietal lobule in response to a given sub-second duration was shown to be maximally reduced by recent exposure to the same duration and progressively less so when the difference in duration between the two repetitions increased (Hayashi et al., 2015). Conversely, motion adaptation shows a very narrow spatial tuning (Ayhan et al., 2009) and no interocular transfer (Bruno et al., 2010) - but see Burr et al. (Burr et al., 2007) - pointing to an early rather than late brain locus. However, we note that our computational model is able to reproduce both of these effects within a single framework, using a common mechanism of adaptation.

Recent neuroimaging studies discovered the existence of chronotopic maps with a topographical organisation in several brain areas, ranging from sensory occipital cortices to motor frontal cortices (Harvey et al., 2020; Protopapa et al., 2019): neuronal populations in these areas show a maximal response to a preferred interval duration, whereas their activity is inhibited by nonpreferred durations. Interestingly, the described timing selectivity seems to depend on the time scale, with a less defined spatial progression of the chronotopic maps for supra-second durations and with variable map sizes depending on the tested duration range. It is clear to see that these maps well capture the characteristics of the ‘duration channels’, which would be affected by duration adaptation (Heron et al., 2012). Even though chronotopic maps were found in visual areas that are involved in motion processing (Harvey et al., 2020), it is far less clear how these brain mechanisms can account for the effects of motion adaptation, like those we report here. One might argue that the adaptation durations we used in this experiment (32 or 8 s) are longer than any of the test durations and, therefore, the duration compression we observed might have resulted from adapting a duration channel, like those we have just described. However, in their study, Heron et al. showed that their duration channels are narrowly tuned around the preferred duration and the effect of adaptation tended to disappear when the difference between adapting and test duration was larger than 1.5 octaves. The same difference in our paradigm is substantially larger than that and, therefore, we should not expect any effect of duration adaptation on our results. Following the same logic, the supra-second duration we used should have shown a bigger compression than the sub-second duration, as it is closer to the adapting duration, but this was not the case in our results.

The idea that any mechanism that can process duration does not need any extra component to process temporal frequency or rate and that, therefore, a single mechanism might exist in our brain to process both (Brighouse et al., 2014; Hartcher-O’Brien et al., 2016) is, to some extent, supported by the results of this study. Temporal frequency, rather than speed, seems to be the key dimension which gets adapted here, as demonstrated by previous observations that adaptation to flicker induces a comparable duration compression (Johnston et al., 2006). It has been recently shown that adaptation to a slow or a fast rate can induce duration overestimation and underestimation, respectively, of empty intervals, i.e., only defined by transient signals at onset and offset (Motala et al., 2020), suggesting that the observed aftereffects do not depend on the content of the test intervals. On the contrary, adaptation to visual motion or flicker biases perceived duration even in the absence of any change in the perceived onset and offset of the test intervals (Gulhan & Ayhan, 2019; Johnston et al., 2006), implying that the content of an interval is taken into account when duration is processed, as hypothesized by some models of duration perception (Johnston, 2010, 2014; Roseboom et al., 2019). The observation that intervals with equal length containing stimuli moving at different speed, or with different speed temporal profiles, are perceived to have different durations (Binetti et al., 2012; Bruno et al., 2015; Bruno et al., 2022; Kanai et al., 2006) also points to a central role of interval content in duration perception. Comparing the actual and subjective durations of intervals containing sinusoidal gratings drifting at a constant speed with that of intervals containing accelerating gratings with the same average speed, in an fMRI study, Binetti et al. observed two separate subsets of activated areas, one including early visual areas for objective durations and one including more anterior areas and the cerebellum for subjective durations (Binetti et al., 2020). This is further evidence that duration perception is achieved by a network of distributed mechanisms with different levels of involvement in the extraction and estimation of duration information from a sensory signal.

Our results also showed that the bias in perceived duration after adaptation could not be ascribed to a reduced ability to discriminate between duration and that, on the contrary, duration discrimination thresholds lowered after adaptation (especially for 600 ms), indicating an enhanced sensitivity. This seems to go against the intuition that adaptation might have increased the internal noise and, therefore, flattened the psychometric functions. As a matter of fact, our finding is in line with other vision studies that showed better discrimination after adaptation to contrast (Abbonizio et al., 2002), orientation (Clifford et al., 2001) or face identity (Oruc & Barton, 2011). Speed adaptation was found to induce subsequent changes in perceived speed accompanied by an increase in speed discriminability (Bex et al., 1999; Clifford & Wenderoth, 1999), the same pattern we observed here for duration. In our case, the changes in apparent speed after adaptation were minimized by the use of a ‘mixed’ adaptor, which alternated between two frequencies (5 and 20 ° /s) known to induce opposite speed aftereffects (i.e., subjective increase and decrease, respectively), which were shown to cancel each other (Ayhan et al., 2011; Ayhan et al., 2009; Bruno et al., 2010). The fact that, even under these conditions, the duration bias did not disappear, is a further demonstration that changes in perceived duration after adaptation are dissociable from changes in perceived speed. To our knowledge, no studies reported any difference in perceived speed between stimuli embedded in sub- versus supra-second intervals or a different effect of 5 and 20 ° /s adaptation on a 10 ° /s test embedded in a supra-second interval.

In conclusion, our results support the idea that the processing of duration information contained in a sensory signal is achieved by multiple mechanisms, which are independently recruited according to the duration range of the intervals to estimate. The advantage of having several different mechanisms, which might appear to represent time in a redundant fashion, is twofold: first, if necessary, they allow us to deal with duration information from multiple intervals simultaneously and, second, as we have different spatial brain maps to guide perception and action, we can benefit from having different mechanisms specialized for different purposes and functions. Adaptation to visual motion, for its well defined spatial and temporal characteristics and for the robustness of the effects it induces on subjective duration, is an ideal tool to investigate the different contributions of cortical and subcortical components to the representation and experience of interval duration at different time scales.

## Supporting information

Electronic Supplementary Material

## Acknowledgements

We would like to acknowledge the support of the Wellcome Trust to this research (213616/Z/18/Z grant awarded to A.B.).

## Data Availability

Experimental and analysis code, and raw and processed data presented in this manuscript were made available here: https://osf.io/ps397/

