## Supplementary material for "A purely visual adaptation to motion can differentiate between perceptual timing and interval timing": Electronic Supplementary Material

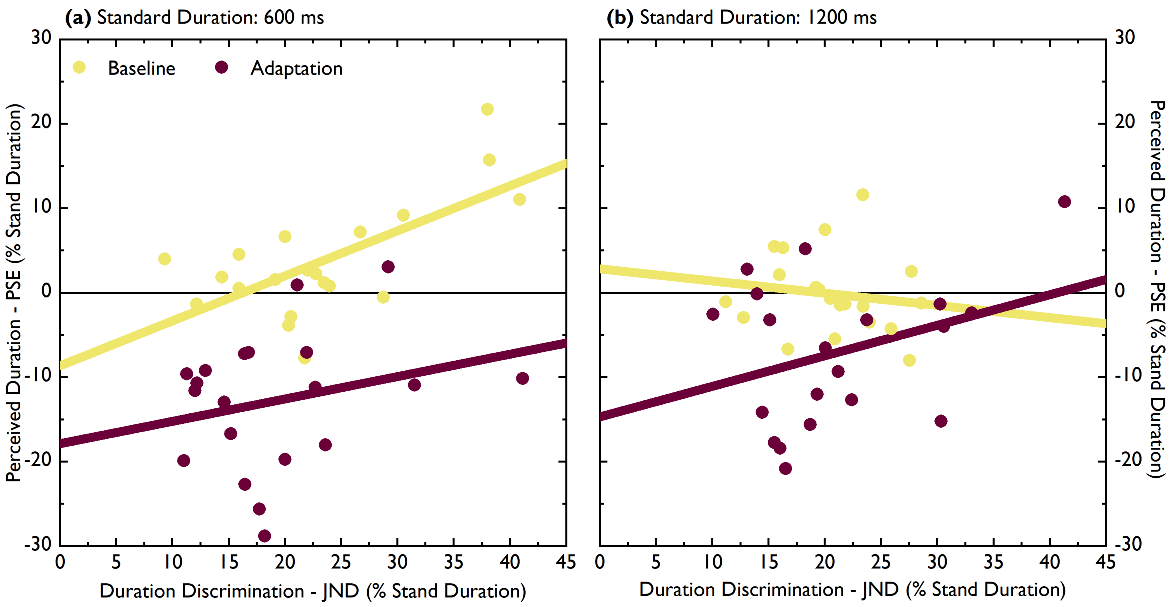


**Figure S1**. The linear fits between the individual PSEs and the individual JNDs for the baseline (yellow) and adaptation (dark red circles and lines) conditions, expressed as percentage changes relative to the standard duration, are plotted for the 600 ms standard duration (panel (a)) and for the 1200 ms standard duration (panel (b)).
